# FEATS: Feature selection based clustering of single-cell RNA-seq data

**DOI:** 10.1101/2020.07.13.200485

**Authors:** Edwin Vans, Ashwini Patil, Alok Sharma

## Abstract

Advances in next-generation sequencing (NGS) have made it possible to carry out transcriptomic studies at single-cell resolution and generate vast amounts of single-cell RNA-seq data rapidly. Thus, tools to analyze this data need to evolve as well to improve accuracy and efficiency. We present FEATS, a python software package that performs clustering on single-cell RNA-seq data. FEATS is capable of performing multiple tasks such as estimating the number of clusters, conducting outlier detection, and integrating data from various experiments. We develop a univariate feature selection based approach for clustering, which involves the selection of top informative features to improve clustering performance. This is motivated by the fact that cell types are often manually determined using the expression of only a few known marker genes. On a variety of single-cell RNA-seq datasets, FEATS gives superior performance compared to the current tools, in terms of adjusted rand index (ARI) and estimating the number of clusters. In addition to cluster estimation, FEATS also performs outlier detection and data integration while giving an excellent computational performance. Thus, FEATS is a comprehensive clustering tool capable of addressing the challenges during the clustering of single-cell RNA-seq data. The installation instructions and documentation of FEATS is available at https://edwinv87.github.io/feats/.

## Introduction

Single-cell RNA-sequencing (RNA-seq) quantifies the gene expression profile of the whole transcriptome of individual cells^1^. Analysis of single-cell RNA-seq data can provide numerous insights such as characterization of cells into sub-types to reveal heterogeneity^2^ and the detection of rare cell types^3^. Advances in sequencing technology have enabled labs to rapidly sequence up to millions of individual cells from different species using varied protocols, library preparation steps, and sequencing machines. Thus, the major challenge is the computational analysis of the enormous amount of data generated by single-cell experiments^4^.

Effective analysis of single-cell RNA-seq data requires computational tools that can accurately perform multiple tasks such as clustering, outlier detection, and integration of multiple datasets. Through clustering, cells can be characterized into types and sub-types. Estimating the number of clusters, and hence the number of cell types, in a dataset is usually performed manually. Selecting too many or too few clusters can result in incorrect cell type assignments. In contrast, outlier detection involves automatic detection of rare or damaged cells that are biologically unrelated to any clusters (but are assigned into one of the groups by clustering) and are located sufficiently far from the central density of the data. Batch effects in the data is a condition where the expression of genes for the same cell types differs across different batches or experiments. It is a technical variation that arises when data is generated in different laboratories, following different protocols, with various library preparation steps or sequencing machines. Such differences if not removed can lead to incorrect biological conclusions. Thus, batch correction is necessary during the integration of multiple datasets.

Several tools address each of these issues independently^5–7^. However, using these tools together is challenging. Here we present a python software package called FEATS, for clustering of single-cell RNA-seq data and estimating the number of groups in the data. FEATS also facilitates the detection of rare cells, through the computation of outliers and performs batch correction and integration of data from multiple experiments.

## Results

### Clustering and estimating the number of clusters

To cluster single-cell RNA-seq data, we use a feature selection based approach^8^. We briefly describe the FEATS clustering pipeline (Fig 1a, Supplementary Note 1). The input is the gene expression count matrix *X* which is a *d* × *n* matrix where *d* represents features (genes), and *n* represents samples (cells). The output is *k* clusters of the input dataset, where *k* is either a known parameter or is estimated by the algorithm. To select features, we first filter genes that are lowly or highly expressed (Methods). This reduces the number of genes to *d*_1_ (Fig 1a). We then normalize the gene expression matrix and perform analysis of variance (ANOVA) on the normalized data (Methods). Essential features, or genes, are selected using ANOVA F-value, i.e., the higher the F-value, the more important the feature. ANOVA can detect features whose expressions vary significantly across different cell types (Supplementary Figure 1). The motivation for this approach is that before complex computational tools were available, cell types were determined manually by observing the gene expression of specific genes or genetic markers. Here, by using univariate feature selection, we choose features that are informative enough to characterize cell types.

**Figure 1.**
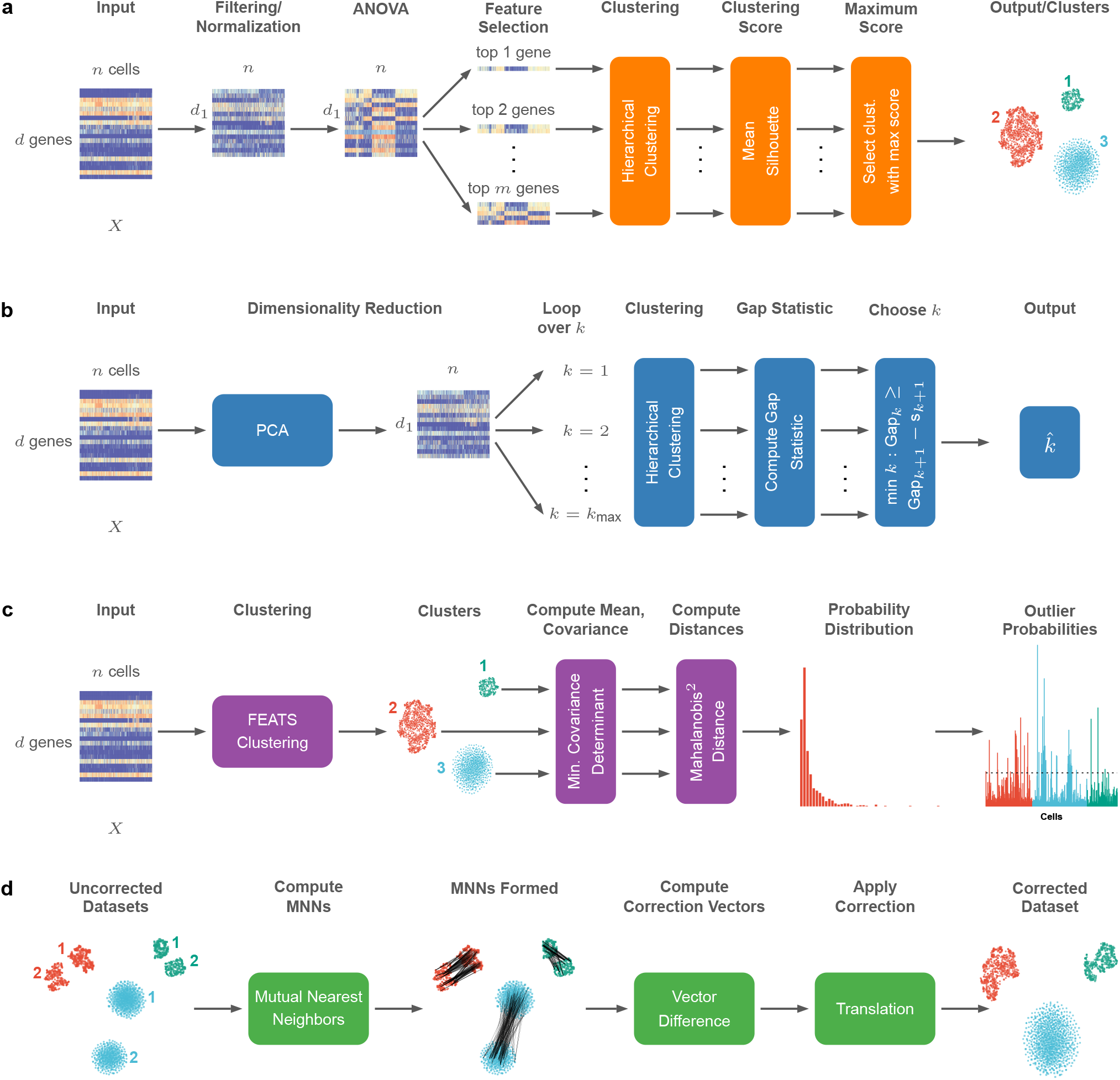
The proposed FEATS clustering, outlier detection, and batch integration schemes. (a) A block diagram showing the computational stages in the proposed FEATS clustering pipeline. Heatmaps are used to represent the gene expression matrix after various stages of the pipeline. After the gene filtering step, the number of genes remaining is *d*_1_. The ANOVA step includes computing temporary clusters, performing ANOVA, and sorting the genes according to the F-value to get the final clusters. (b) The gap statistic approach of FEATS to estimate the number of clusters in the dataset. PCA is used to reduce the dimensionality and for *k* = 1, 2, …, *k*_max_ clustering is performed and the gap statistic is computed. The estimated number of clusters is chosen as the minimum *k* value that satisfies Gap(*k*) ≥ Gap(*k* + 1) − *s*_*k*+1_ (c) The outlier detection pipeline of FEATS. The clustering step here refers to performing FEATS clustering outlined in (a). Once clustering is performed, the mean and covariance of the samples in each cluster are computed using the Minimum Covariance Determinant (MCD) algorithm. The squared Mahalanobis distance is computed for all the cells using their respective cluster mean and covariance. (d) An overview of the Mutual Nearest Neighbour (MNN) batch correction technique used to correct batch effects in the data. In this diagram, two batches (1 and 2) with the same cell types (red, blue, green) are shown before applying batch correction. The MNN connections are shown as black lines connecting the mutual nearest neighbors formed between the two batches. After batch correction similar cell types appear together.

The next step in the clustering pipeline is to determine the optimal number of informative features, or genes, to choose for clustering. To do this, we use a forward selection method in which we iteratively choose top *q* features, where *q* = 1, 2, …, *m*. Here, *m* is the maximum number of features. We choose *m* as 5% of *d* to avoid noisy features being selected and to keep computational time low. Once the top *q* features are selected in each iteration, hierarchical clustering is performed and a clustering score is computed. The clustering score is used to assess the quality of clusters formed for every top *q* features selected. Here we utilize the silhouette coefficient (Methods). The silhouette coefficient is a measure of how tightly samples are grouped into clusters. The clustering score is computed as the mean silhouette coefficient of all the samples (Supplementary Figure 2). The final output of the algorithm is the groupings which achieve the maximum clustering score.

We have selected nine single-cell RNA-seq datasets that are publicly available (Fig 2a), to compare and benchmark the FEATS clustering method against other techniques. These datasets contain a variety of cells from human and mouse, such as cells from mouse embryo development at various stages (Biase^9^, Goolam^10^, Deng^11^, Fan^12^), mouse embryonic stem cells under three different culture conditions (Kolodziejczyk^13^), mouse lung tissues (Treutlein^14^), human embryo development at various stages (Yan^15^), human brain cancer tissues from five patients (Patel^16^), and diverse human tissues containing blood cells, pluripotent cells, dermal/epidermal cells and neural cells (Pollen^17^). The performance of our clustering approach on these datasets was computed using the adjusted rand index (ARI) metric (Methods). We have considered six other clustering methods to compare with our results. These are SC3^5^, Seurat^18–20^, pcaRdeuce^21^, SIMLR^22^, SINCERA^23^ and TSCAN^24^. The average performance of our method was better than the rest on the nine datasets (Fig 2b, Supplementary Table 1). We have also confirmed via Wilcoxon signed-rank test with p-value < 0.01 that our method performs better than the other methods with only one exception.

**Figure 2.**
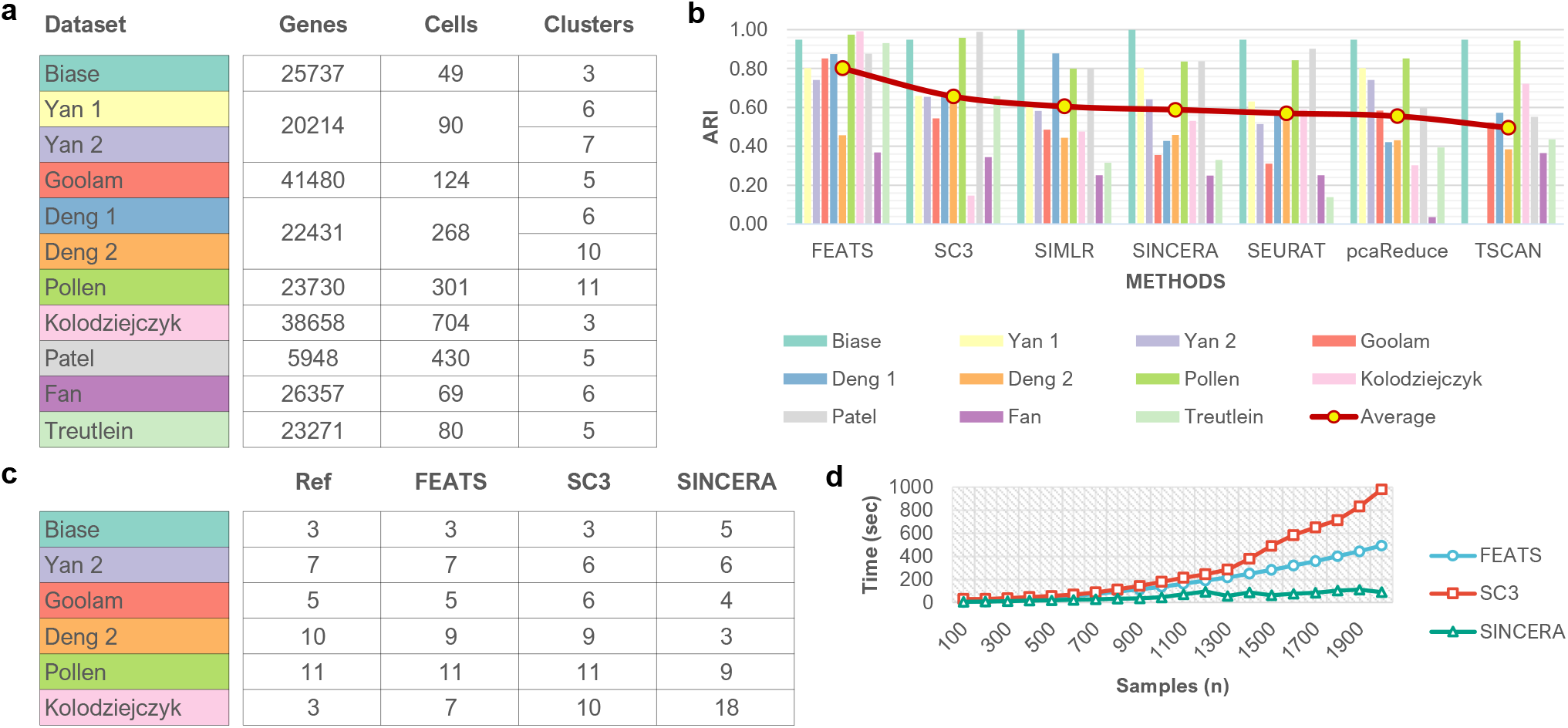
(a) Summary of datasets used to assess the FEATS clustering method. Each dataset is represented using a different color. The cell types in the Yan et al. and the Deng et al. datasets have two distinct hierarchies. (b) Bar plot shows the clustering metric ARI of FEATS clustering compared to other methods on various datasets. The clustering metric ARI is used to compare the results. The circle marker on the bar plot shows the average ARI of the method on all the datasets. For the TSCAN method, no results were obtained for the Yan 1 and Yan 2 datasets. (c) Results of estimating *k* on datasets with high confidence cell labels. The Ref column is the number of clusters as determined by the authors in the original study. (d) CPU time in seconds versus the number of samples (*n*) for the FEATS, SC3 and SINCERA clustering methods.

FEATS is also capable of estimating the number of clusters, *k*, in the data if this is not known by the user (Fig 1b). To do this, we have employed the gap statistic originally proposed by Tibshirani et al.^25^ (Methods, Supplementary Note 2). On datasets with high confidence cell labels (datasets which contain cells from different stages, conditions, or lines), FEATS can estimate the number of clusters more accurately (Fig 2c) when compared to other methods that are capable of determining the number of clusters. FEATS correctly estimated the number of groupings in 4 out of the 6 datasets, whereas, SC3 correctly estimated the number of clusters in 2 out of the 6 datasets.

Additionally, we have evaluated the computational speed of the FEATS clustering method and compared it with the SC3 and SINCERA clustering approaches (Fig 2d). The evaluation was done on a workstation with Intel Core i7-7700HQ @ 2.8GHz (4 cores) CPU with 16GB RAM and 64-bit Windows 10 operating system. To do this evaluation, a simulated dataset with 2000 samples was used. The number of genes was kept constant at around ~ 9000 and samples were randomly selected from the dataset starting from 100 samples in steps of 100 up to 2000 samples. To cluster a single-cell dataset with 2000 cells, SINCERA took around ~ 1.5 minutes, FEATS took around ~ 8 minutes and SC3 took twice the amount of time as FEATS of around ~ 16 minutes. FEATS clustering method showed better computational performance compared to the more accurate SC3 clustering method.

### Outlier detection

To detect outliers, we compute the squared Mahalanobis distance (Methods, Supplementary Note 3) for all cells in each cluster after performing FEATS clustering on the data (Fig 1c). To compute the squared Mahalanobis distance, robust mean and covariance matrix is used so that the outlying cells are not used in the computation of these parameters (Methods). For clusters with a minimal number of cells where robust covariance matrix cannot be computed, FEATS generates a warning and skips computing the outliers for those clusters. This is rarely the case as most datasets available presently have a large number of cells. The next step in our outlier detection scheme is to determine a probability distribution of the squared Mahalanobis distances. The distribution of the squared Mahalanobis distances follows a 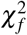 distribution (Supplementary Figure 4) with the degree of freedom *f* equal to the dimensionality of the cells. We use the *χ*^2^ distribution to compute a p-value for each cell. The p-value is the probability of examining a cell with a more extreme squared Mahalanobis distance. We also compute an outlier score for each cell and use a threshold to determine if cells are outliers (Methods, Supplementary Figure 4). We have tested this approach on the mouse intestine data^26^ and compared the results with the popular RaceID^6^ method. FEATS was able to detect only one outlier using a p-value threshold of 10^−4^, while RaceID identified 8 outliers. The sample identified as an outlier using FEATS was also identified as an outlier by RaceID. Different number of outliers identified by FEATS and RaceID was mainly due to different clustering approaches of the two methods.

### Batch correction and integration

We have integrated the mutual nearest neighbor (MNN) algorithm^7^ in FEATS to correct batch effects in the input dataset (Fig 1d). The MNN algorithm has three main assumptions. The first is that at least one cell type is common between the two batches. The second is, that the batch effect is orthogonal to the biological subspace of the two batches. The third assumption is that the variation in the batch effect is much smaller than the biological variation. The MNN batch correction algorithm works by first identifying MNN pair cells between the two batches that are to be merged; a reference batch and a target batch. The batch effect vector is computed as the cell differences of the identified MNN pairs. A batch effect vector for all the cells in the target batch is calculated using the weighted average (via Gaussian kernel weights) of MNN pair specific batch effect vectors. This allows the batch effect vectors to vary smoothly across all the cells. To maintain the orthogonality assumption, the non-orthogonal components from the batch vector are removed by computing a span of both the batches. To correct the batch, the batch effect vector is subtracted from the target batch.

To show the effectiveness of using FEATS to correct and integrate samples from different batches, we have compared the results to another recent method called Scanorama^27^. Firstly, we have generated artificial data in two batches consisting of three different cell types (Fig 3a). All the cell types in the two batches are similar. We have generated t-SNE^28^ plots to show correction and integration results using Scanorama (Fig 3b) and FEATS (Fig 3c). Additionally, we have also compared FEATS and Scanorama on real data. We have taken cells from 293T, Jurkat, and a 50:50 mixture of 293T and Jurkat cells^29^ from the commercial platform 10X Genomics. Here, the 50:50 mixture of 293T and Jurkat cells are not properly aligned with their respective cells (Fig 3d). FEATS (Fig 3f) can correct and integrate the datasets so that the cells are properly aligned with an accuracy similar to Scanorama (Fig 3e).

**Figure 3.**
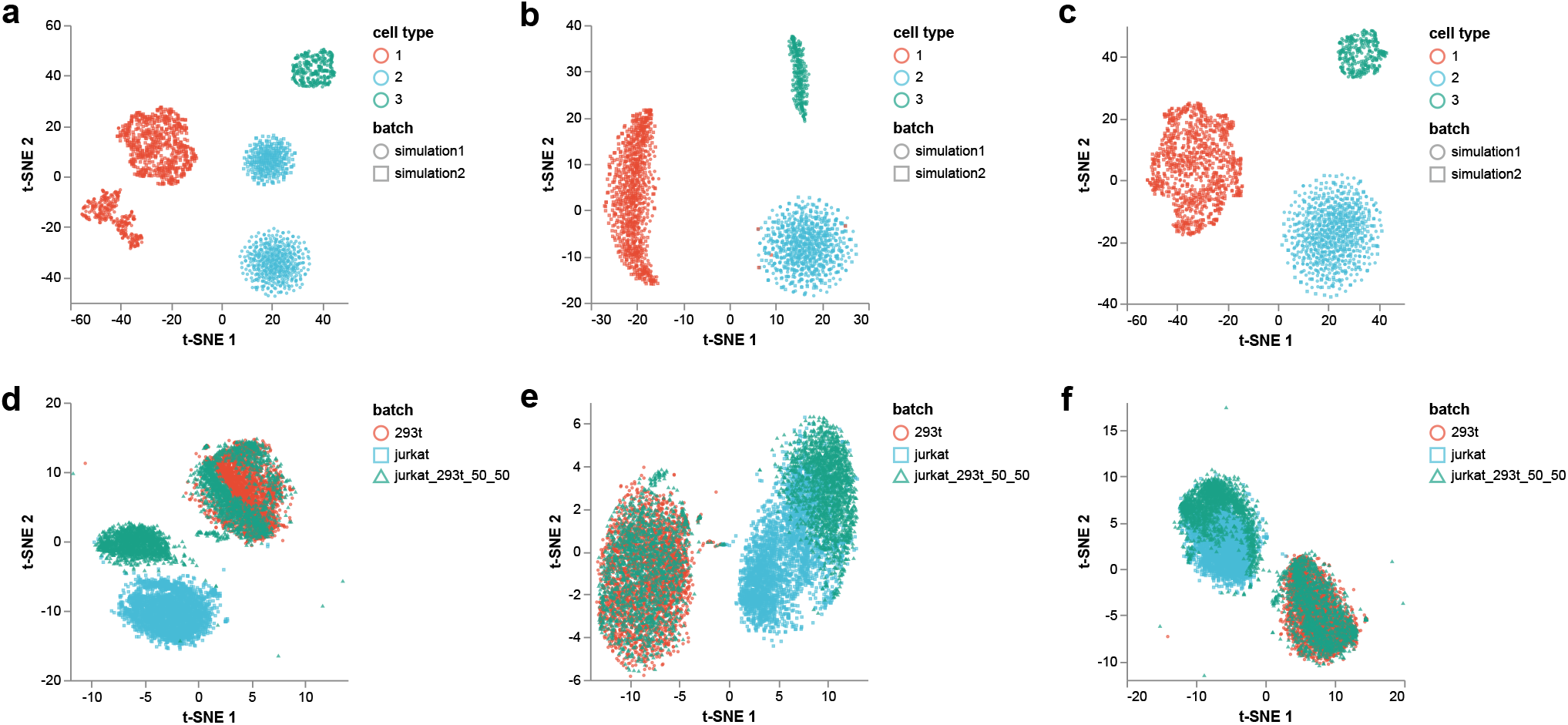
t-SNE plots are illustrating the effects of batch correction and integration on two sets of data (simulated data and real dataset). (a) Uncorrected simulation data consisting of two batches and three cell types. Different batches are shown using different markers and marker colors are by cell types. (b) Simulation data corrected using Scanorama. (c) Simulation data corrected using FEATS. (d) Uncorrected data from three batches 293t, Jurkat, and a 50:50 mixture of 293t and Jurkat cells. The three batches, representing 293t and Jurkat cell types, are shown using different colors and markers. (e) The t-SNE plot showing datasets corrected using Scanorama. (f) The t-SNE plot showing datasets corrected using FEATS. After correction, both (e) and (f) show that the cells are aligned.

## Discussion

The FEATS package provides the user with functions to perform several types of downstream analysis on single-cell RNA-seq datasets. It is capable of performing clustering, estimating the number of clusters, performing outlier detection, and performing batch correction and integration of multiple datasets. Our results show that FEATS gives a superior clustering performance on a range of datasets in terms of ARI when compared to other recently contributed methods. On average FEATS shows a 22% improvement over the SC3. The ability of FEATS to estimate the number of clusters, *k*, is also remarkable. Additionally, FEATS clustering is less complex and takes significantly less computation time when compared to other accurate but more complex clustering methods such as SC3.

Although FEATS clustering gives superior computational performance compared to SC3, the running time is still polynomial. This means that to cluster single-cell datasets with hundreds of thousands of cells on workstations with limited computational resources will take a considerable amount of time. For very large datasets (> 100, 000 cells), we suggest applying a hybrid approach. In the hybrid approach, a fraction of cells can be uniformly sub-sampled from the dataset, and FEATS clustering can be used to cluster the data. For the remaining cells, a supervised approach such as DeepInsight^30^ can be taken in which case the clustered cells will become the training data used to assign cells to clusters.

Compared to other methods, the FEATS clustering method has fewer parameters requiring user selection. One of the parameters is *k*, which is the number of clusters in the data. For this parameter, users can pass an integer, an array of integers representing possible *k* values or the string ‘gap’ which is the default. If an array of integers is passed, FEATS will perform clustering using all the values in the array and store the results. If the string ‘gap’ is passed, FEATS will use gap statistic to compute *k* using *k* from 1, 2, …, *k*_max_. Users can also choose *k*_max_ which by default is 20. The second parameter is *m*, which is the maximum number of top features to select. For this parameter, the user can select an integer between 1 and *d*_1_. Values closer to *d*_1_ will result in more computation time and less important features being selected, whereas values closer to 1 might not result in accurate clustering.

For outlier detection in FEATS, users can provide a reduced dimension which by default is 2. FEATS will then use PCA to reduce the dimensionality before performing outlier detection. Another user-selectable parameter is the p-value threshold for outlier classification, which by default is 10*−4*. To perform batch correction and integration, FEATS first analyzes datasets and determines common genes among the datasets. If no common genes are found an error message is generated. If common genes are located between datasets, FEATS uses these genes to perform integration and correction using the MNN method.

FEATS provides two gene filtering approaches. One approach is where poorly or highly expressed genes can be filtered by specifying filter parameters (Methods). FEATS can also perform gene filtering using highly variable genes (HVGs) by computing the dispersion for each gene. Here, users will need to provide the number of HVGs to select. For normalizing the gene expressions, FEATS provides several methods such as z-score normalization, *L*^2^ normalization, and cosine normalization.

In conclusion, FEATS is a comprehensive single-cell data analysis tool that combines a fast clustering algorithm that is superior to existing methods with outlier detection and batch correction showing a performance similar to currently used tools.

## Methods

### Gene filtering and normalization

As a preprocessing step, we take the gene expression counts matrix where rows represent genes and columns represent cells and apply log transformation after adding a pseudo-count of 1. Thus, we get *X* = log_2_(counts + 1). A gene filter is applied which rejects highly and lowly expressed genes. The gene filter removes genes expressed (with expression value > 0) in less than *r*_min_ number of cells and genes expressed in greater than *r*_max_ number of the cells. For our clustering experiments we choose *r*_min_ as 10% of *n* and *r*_max_ as 90% of *n*, since ubiquitous and rare genes do not help much in clustering. The filtering criterion also reduces the number of genes in the dataset, increasing the computational speed of the method. Finally, the features are normalized using the z-score normalization.

### ANOVA

The FEATS clustering approach performs ANOVA on the filtered and normalized gene expression input matrix. To do this, temporary clustering is computed using the agglomerative hierarchical clustering employing the Ward linkage criterion. The temporary grouping of the input may not be very accurate (Supplementary Figure 1). ANOVA is performed using temporary cluster labels. In statistical hypothesis testing, ANOVA is used to test the null hypothesis that the mean of a variable, such as the expression of a gene, is the same for all the groups; *H*_0_ : *μ*_1_ = *μ*_2_ = … = *μ*_*k*_. Using ANOVA, an F-statistic is computed from which a p-value corresponding to each of the features can be obtained. Usually, p-values are used to reject the null hypothesis, for example, if p is less than 0.05 (at 5% level of significance). However, we use F-values for our feature selection approach. An F-value is a ratio of between-class to within-class variances. The goal is to select features that maximize this ratio. Each feature will be assigned an F-value and the higher the F-value, the more critical the corresponding feature is. FEATS uses the popular scikit-learn^31^ implementation of ANOVA functions.

### Gap Statistic

To estimate the number of clusters in the data, FEATS uses the gap statistic. The gap statistic method compares the within-cluster variance *W*_*k*_ for different *k* values with its expected value under a null reference distribution of the data, i.e., a uniform random distribution. The estimate of the optimal number of clusters 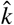 is the value for which gap statistic is the maximum. The gap statistic is defined as follows

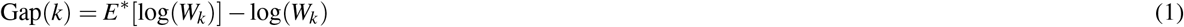

To compute the gap statistic, we first perform PCA on the dataset to reduce the dimensionality (Supplementary Note 2). In addition to reducing computational complexity, this reduces noise in the data. The reduced dimension is selected such that the explained variance is 99% or more. We generate the reference datasets using a bounding box in the reduced space. To obtain the estimate of *E*\*[log(*W*_*k*_)] we compute the average of *B* copies of 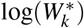. Thus, we have

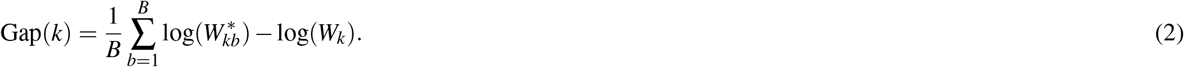

Here, we use *B* = 500 to strike a balance between accuracy and speed. To choose the optimal value for *k*, we compute the standard deviation sd(*k*) as

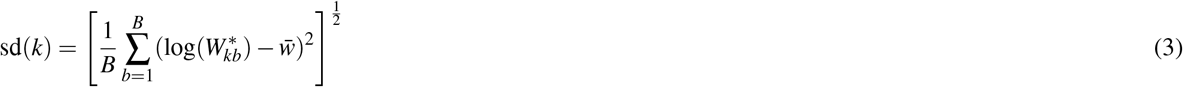

where 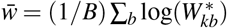, and we also define *s*_*k*_ as

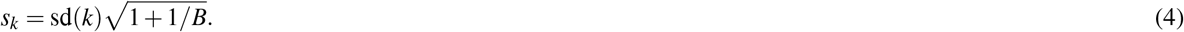

Finally, the optimal number of clusters 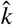 is selected as the smallest *k* such that Gap(*k*) ≥ Gap(*k* + 1) − *s*_*k*+1_ (Supplementary Figure 3). We use hierarchical clustering with Ward criterion to cluster the dataset and the null reference datasets for computation of gap statistic. FEATS computes gap statistic for *k* from 1, 2, …, *k*_max_, where *k*_max_ is an integer which can be defined by the user.

### Outlier Probabilities

To perform outlier detection on datasets, we first cluster the data using the FEATS clustering method and reduce the dimensionality of the cells to 2-dimensions using PCA (Supplementary Note 3). We then compute the squared Mahalanobis distance of all the samples within each cluster using the cluster mean and covariance. The squared Mahalanobis distance is defined as

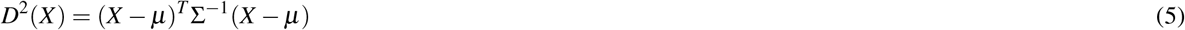

where *μ* is the sample mean and Σ is the sample covariance matrix. Here, the shape of the data is taken into account through the sample covariance matrix. However, outliers are masked when computing the standard sample mean and covariance matrix. A robust way of computing the mean and the covariance matrix is needed so that the outlying samples are not used in the computation. We use Rousseeuw’s minimum covariance determinant (MCD) algorithm^32^ as a robust estimator for the mean and covariance of multivariate data with outliers. The MCD algorithm is used to find *h* samples out of *n* whose covariance matrix has the lowest determinant. We use the efficient version of MCD^33^ implemented in the Python scikit-learn machine learning package to compute the robust mean and covariance for each cluster.

To compute the outlier probabilities, the distribution of the Mahalanobis squared distance is considered. The Mahalanobis squared distance is 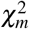 distributed, where *m*, which is the reduced dimensionality of *x*_*i*_ and is also the degree of freedom of the *χ*^2^ distribution. A p-value for each sample is computed using this distribution and the squared Mahalanobis distance. We also compute an outlier score for each sample as −log_10_(p-value). Finally, the cells with a p-value less than the threshold 10^−4^ are classified as possible outliers.

### Silhouette Score

The clustering score is calculated as the mean of the individual silhouette coefficient of the samples. The silhouette coefficient for each sample is computed as follows

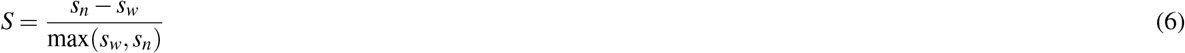

where *s*_*w*_ is the mean of within-cluster distance, and *s*_*n*_ is the mean nearest cluster distance for the sample.

### Adjusted Rand Index

We use the ARI metric to compare two different clustering runs. The ARI metric is defined as follows. Given a set of *n* samples in the data and two groups of these samples, the overlap between the two groupings can be summarized in a contingency table, where each entry *n*_*ij*_ represents the number of samples in common between the *i*-th group of the first clustering and the *j*-th group of the second clustering. The adjusted rand index is computed as follows

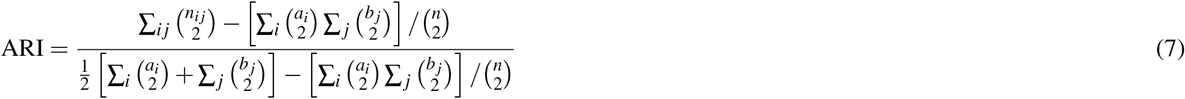

where *a*_*i*_ and *b*_*j*_ are the sums of rows and columns of the contingency table respectively. The ARI values are in the range [−1, 1]. A value of 1 indicates perfect grouping. A value of 0 indicates a random assignment of samples to groups, and negative values indicate wrong cluster assignments. For all our clustering experiments, we have taken the median ARI of 100 trial runs as some clustering methods were not giving stable results.

### t-SNE Plots

To generate the t-SNE plots, we have used the scikit-learn implementation of the *TSNE* function to reduce the data to two dimensions. We have used the following parameters of the *TSNE* function to generate two-dimensional mappings. The perplexity parameter, learning rate and the number of iterations is set to 600, 200 and 400 respectively, and we have selected PCA as the initialization of the embeddings.

## Data Availability

All the datasets used in this paper are publicly available. The datasets Biase, Yan, Goolam, Deng, Pollen, Kolodziejczyk, Patel, Fan and Treutlein were downloaded in RDS format from https://hemberg-lab.github.io/scRNA.seq.datasets/. The counts/normalized counts and column/row data was extracted and stored in comma separated values (CSV) file format for access using Python. The Jurkat and 293T cell datasets were downloaded from http://cb.csail.mit.edu/cb/scanorama/.

## Code Availability

FEATS is available as a Python package, and the latest version can be installed from the Python package index (PYPI) using the command `pip install feats`. Source distribution can be obtained from our GitHub repository https://github.com/edwinv87/feats/tree/master/dist. The Python and R scripts used to produce the results and figures in the paper can also be obtained from our GitHub repository https://github.com/edwinv87/feats/tree/master/scripts.

To run experiments on existing methods, we have installed their software packages as follows:

- The latest Seurat R package was installed using the instructions on https://satijalab.org/seurat/ on May 20, 2020.
- The latest SC3 R package was installed from Bioconductor on May 20, 2020.
- The latest SIMLR R package was installed from Bioconductor on May 20, 2020.
- The pcaReduce clustering package was downloaded from GitHub https://github.com/JustinaZ/pcaReduce on May 21, 2020 and installed using the instructions in the read-me file.
- The SINCERA package was installed on May 21, 2020 by following the instructions on their GitHub page https://github.com/xu-lab/SINCERA.
- The latest TSCAN R package was installed from Bioconductor on May 21, 2020.
- The Scanorama Python package was installed from PYPI on May 21, 2020.
- The RaceID source code was downloaded from https://github.com/dgrun/RaceID on June 18, 2020.

## Author contributions

E.V. developed the method, wrote the first draft of the manuscript and prepared the figures, A.P. and A.S., perceived, supervised and contributed in paper writing.

