## Supplementary Figures and Notes for "FEATS: Feature selection based clustering of single-cell RNA-seq data"

July 13, 2020

#### Supplementary Figures

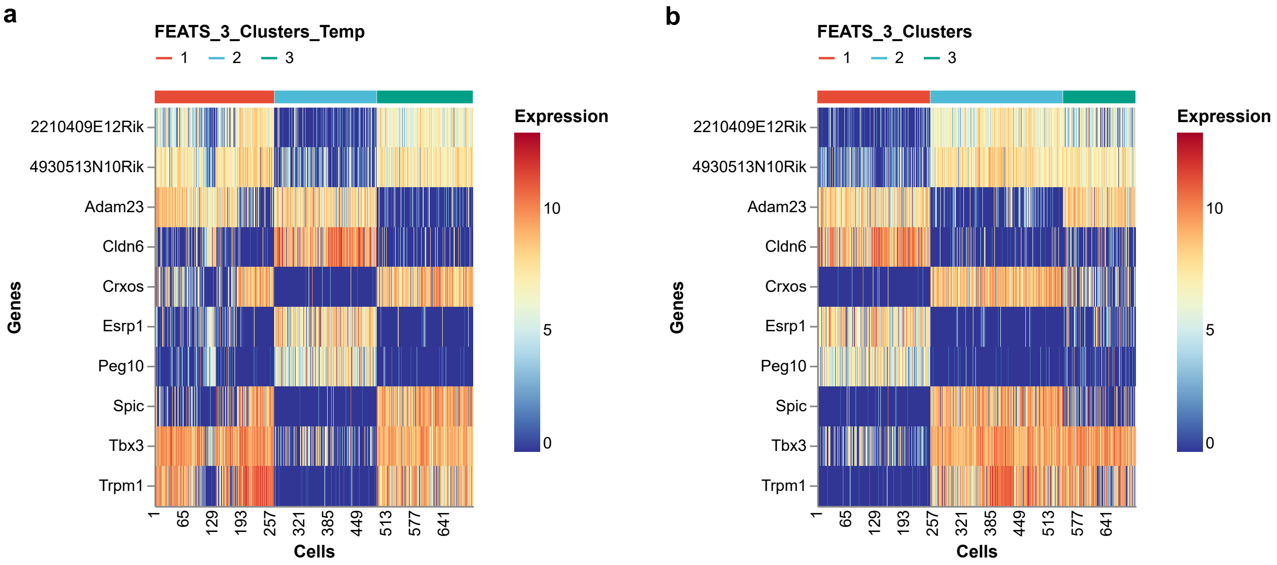


Supplementary Figure 1: Heatmaps showing log-transformed expressions of the top 10 genes after performing ANOVA in an example (Kolodziejczyk et al.) dataset. Here, temporary clusters are used to perform ANOVA. (a) The cells are sorted according to the temporary clusters. It is seen that ANOVA can find genes whose expressions discriminate between temporary classes. It is also visible that the temporary clusterings are not accurate. The gene expressions of some cells in cluster 1 are different which suggests that these cells do not belong in cluster 1. (b) The cells are sorted according to the final clustering output of FEATS. FEATS is able to correct and improve the clusterings so that the cells which are mis-classified in the temporary groupings are corrected.

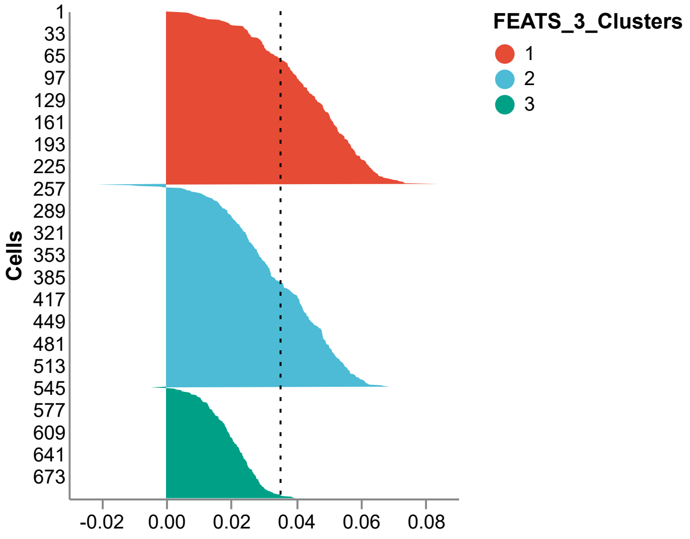


Supplementary Figure 2: Silhouette plot for the Kolodziejczyk et al. dataset after performing clustering. The silhouette plot shows the silhouette index for each sample. Here the silhouette index within each cluster is sorted and different colors are used to show the clusters formed using FEATS. The clustering score is the mean of silhouette coefficient of all the samples. The mean is shown using a black dashed line.

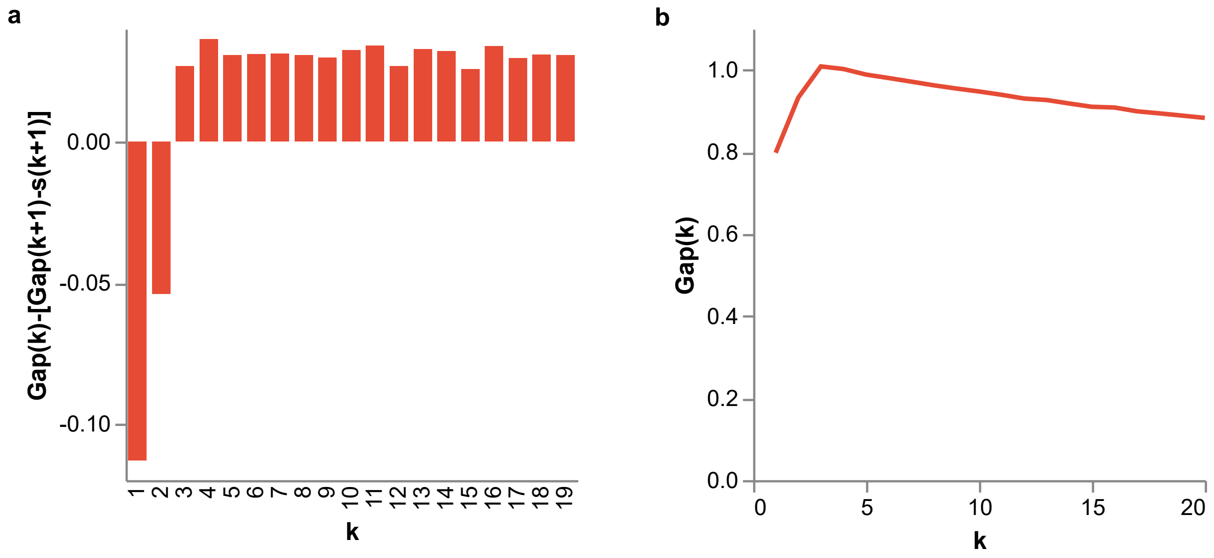


Supplementary Figure 3: Estimation of the number of clusters $k$ for the Biase et al. dataset for $k = 1, 2, ..., 20$ (a) The bar plot shows that $\text{Gap}(k) - (\text{Gap}(k+1) - s_{k+1})$ is positive for the first time when $k = 3$, hence, the algorithm correctly estimates $k$ as 3. This is the criteria used by FEATS to estimate $k$. (b) The plot of the gap statistic versus $k$ also shows that gap is maximum when $k = 3$. However, this may not always be the case for all the datasets.

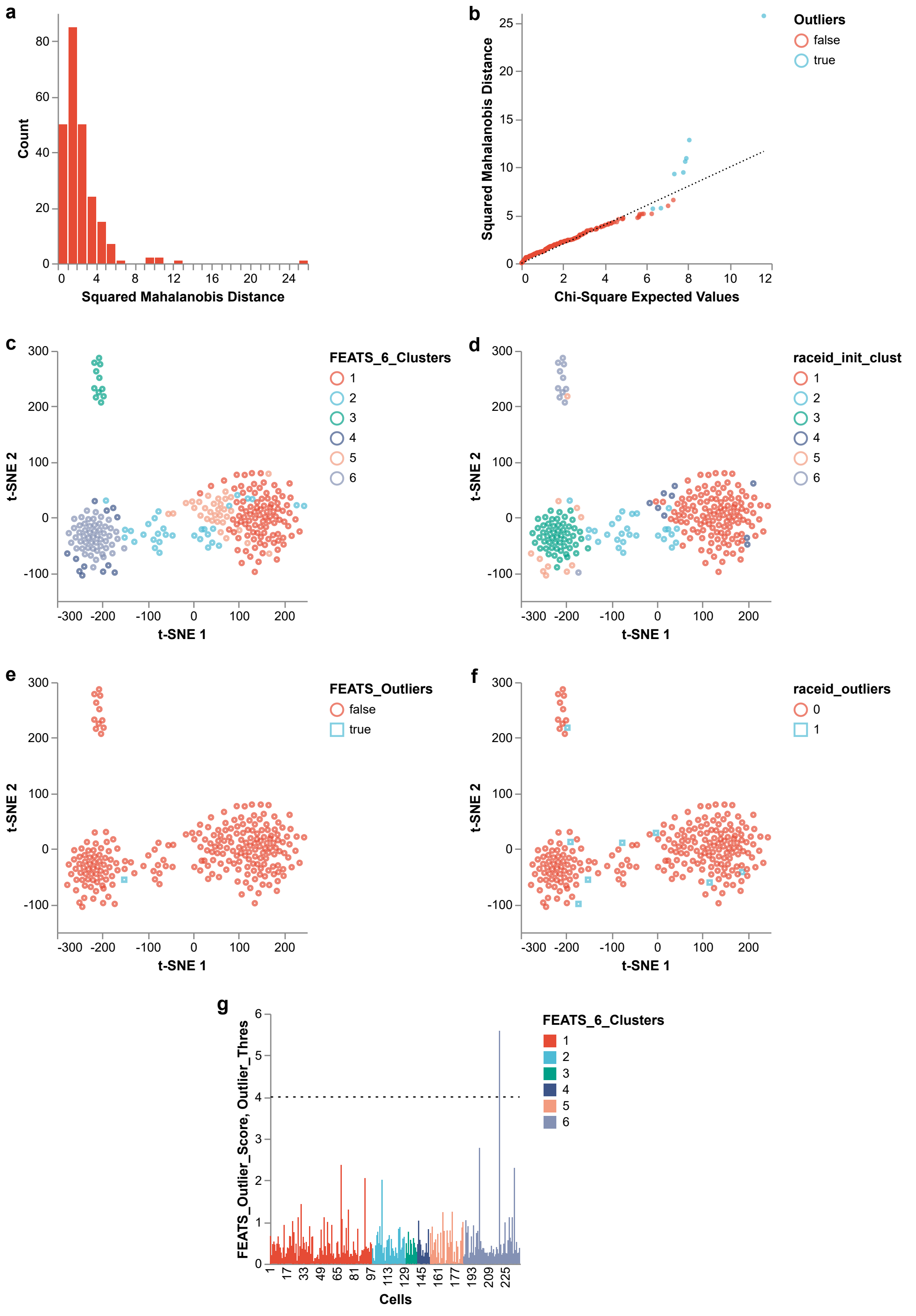


Supplementary Figure 4: Figures for the outlier analysis of the mouse intestine dataset. (a) Histogram showing the distribution of squared Mahalanobis distance. (b) Squared Mahalanobis distance versus sorted $\chi^2$ expected values with degree of freedom 2. The black dotted line represents $y = x$. It shows that the squared Mahalanobis distance follows the $\chi^2$ distribution. Outlier data points here refer to the outlying samples which are not used by the minimum covariance determinant (MCD) algorithm in computation of the robust mean and covariance for each cluster. (c) t-SNE plots showing clusters obtained by the FEATS clustering method and (d) by RaceID. (e) t-SNE plots showing samples that are classified as outliers using FEATS and (f) using RaceID. The blue square data markers are used to represent samples which are outliers. (g) Outlier scores versus samples sorted according to their cluster assignments. The dashed line shows the p-value threshold of $10^{-4}$ used in FEATS to determine outliers. The outlier scores are $-\log_{10}$(p-value)

### Supplementary Notes

#### Supplementary Note 1 - FEATS Clustering Algorithm

The proposed FEATS clustering algorithm is given in Algorithm 1. The inputs to the algorithm are as follows:

- $X$ which is a $d \times n$ gene expression matrix where $d$ is the number of genes (features) and $n$ is the number of cells (samples)
- $k$, where $1 < k < n$ which represents the number of clusters in the data. The user will have to define parameter $k$ here and it can also be estimated by FEATS (see Algorithm 2).
- $m$, where $1 \leq m \leq d_1$, $d_1 \leq d$ which is the maximum number of features to select (default: 5% of $d_1$).

The output is $y$, the cluster labels of the cells in $X$.

**Algorithm 1:** Proposed FEATS Clustering Algorithm

**Input:** $X \in \mathbb{R}^{d \times n}$, $k$ and $m$

**Output:** $y \in \{\omega_1, \omega_2, ..., \omega_n\}$, where $\omega_i \in \{1, 2, ..., k\}$ is the cluster labels for $i = 1, 2, ..., n$

$X_{\text{filt}} \leftarrow \text{gene\_filter}(X, r_{\min}, r_{\max})$, where $X_{\text{filt}} \in \mathbb{R}^{d_1 \times n}$, $d_1 \leq d$, $r_{\min}$ is the minimum and $r_{\max}$ is the maximum number of cells in which genes are expressed with expression value greater than 0.

$X_{\text{norm}} \leftarrow \text{normalize}(X_{\text{filt}}, \text{norm\_type})$, where $\text{norm\_type} \in \{\text{'l2'}, \text{'mean'}, \text{'cosine'}\}$

$y_{\text{temp}} \leftarrow \text{hierarchical\_clustering}(X_{\text{norm}}, k)$

$\text{f\_score} \leftarrow \text{ANOVA}(X_{\text{norm}}, y_{\text{temp}})$

Arrange the genes in $X_{\text{norm}}$ according to the values of f_score

**for** $q = 1$ *to* $m$ **do**

    Choose the top $q$ genes from $X_{\text{norm}}$ to get $X_{\text{red}} \in \mathbb{R}^{q \times n}$

    $y_q \leftarrow \text{hierarchical\_clustering}(X_{\text{red}}, k)$, where $y_q \in \{\omega_1^q, \omega_2^q, ..., \omega_n^q\}$, $\omega_i^q \in \{1, 2, ..., k\}$ for $i = 1, 2, ..., n$

    $\text{cs}_q \leftarrow \text{average}[\text{silhouette\_score}(X, y_q)]$

    Save $\text{cs}_q$, $y_q$

$i \leftarrow \arg\max_q \text{cs}_q$, where $1 \leq i \leq m$

$y \leftarrow y_i$

#### Supplementary Note 2 - Gap Statistic Algorithm

The gap statistic approach of FEATS to estimate the number of clusters in the data is given in Algorithm 2. The inputs are:

- $X$ which is a $d \times n$ gene expression matrix where $d$ is the number of genes (features) and $n$ is the number of cells (samples)
- $k_{\text{max}}$, where $2 \leq k_{\text{max}} \leq n$ the upper limit of the number of clusters (default: 20, with the assumption $n > 20$).
- $B$, a scalar representing the number of reference datasets to generate (default: 500).

The output is $\hat{k}$, the extimate of the number of clusters in $X$.

---

**Algorithm 2:** Gap Statistic Algorithm

---

**Input:** $X \in \mathbb{R}^{d \times n}$, $k_{\text{max}}$ and $B$
**Output:** $\hat{k}$, the estimate of the number of clusters

$X_{\text{red}} \leftarrow \text{PCA}(X, 0.99)$, where $X_{\text{red}} \in \mathbb{R}^{d_1 \times n}$ and $d_1$ which is $< d$ is the number of features remaining after dimentionality reduction using PCA so that 99% of variance is retained.
(x_min, x_max) $\leftarrow$ bounding_box($X_{\text{red}}$), where (x_min, x_max) $\in \mathbb{R}^{d1}$, are the minimum and the maximum gene expression values of $X_{\text{red}}$ in each dimension.
**for** $k = 1$ *to* $k_{max}$ **do**
  $y \leftarrow$ hierarchical_clustering($X_{\text{red}}, k$), where $y \in \{\omega_1, \omega_2, ..., \omega_n\}$, $\omega_i \in \{1, 2, ..., k\}$ for $i = 1, 2, ..., n$
  $W_k \leftarrow \sum_{i=1}^{k} \sum_{x_j \in C_i} \|x_j - \mu_i\|^2$, where $C_i$ represents the $i$-th cluster and $\mu_i$ is the $i$-th cluster mean for $i = 1, 2, ..., k$ and $x_j$ is the $j$-th sample of $X_{\text{red}}$ in the $i$-th cluster.
  Save $W_k$
  **for** $b = 1$ *to* $B$ **do**
    $X_{\text{b}} \leftarrow$ uniform(x_min, x_max), where $X_{\text{b}} \in \mathbb{R}^{d_1 \times n}$ is a matrix of uniform random values within the x_min and x_max bounds.
    $y_b \leftarrow$ hierarchical_clustering($X_{\text{b}}, k$), where $y_b \in \{\omega_1^b, \omega_2^b, ..., \omega_n^b\}$, $\omega_i^b \in \{1, 2, ..., k\}$ for $i = 1, 2, ..., n$
    $W_k^b \leftarrow \sum_{i=1}^{k} \sum_{x_j^b \in C_i^b} \|x_j^b - \mu_i^b\|^2$, where $x_j^b$ is $j$-th sample of $X_b$ in the $i$-th cluster and $\mu_i^b$ is the $i$-th cluster mean.
    Save $W_k^b$
  $\bar{\omega}_k \leftarrow \frac{1}{B} \sum_{b=1}^{B} \log(W_k^b)$
  $\text{Gap}_k \leftarrow \bar{\omega}_k - \log(W_k)$
  $\text{sd} \leftarrow \sqrt{\frac{1}{B} \sum_{b=1}^{B} [\log(W_k^b) - \bar{\omega}_k]^2}$
  $\text{s}_k \leftarrow \text{sd}\sqrt{1 + \frac{1}{B}}$
$\hat{k} \leftarrow$ smallest $k$ such that $\text{Gap}_k \geq \text{Gap}_{k+1} - \text{s}_{k+1}$

---

#### Supplementary Note 3 - FEATS Outlier Detection Scheme

The FEATS outlier detection scheme is given in Algorithm 3. The inputs to the algorithm are as follows

- $X$ which is a $d \times n$ gene expression matrix where $d$ is the number of genes (features) and $n$ is the number of cells (samples).
- The cluster labels $y$, where $y \in \{\omega_1, \omega_2, ..., \omega_n\}$, $\omega_i \in \{1, 2, ..., k\}$ for $i = 1, 2, ..., n$.
- The reduced dimension $d_1$ (delault: 2).
- outlier_thres, the outlier probability threshold (default: $10^{-4}$).

The output is the outlier probabilities (outlier_scores $\in \mathbb{R}^n$) and whether samples are outliers (outliers $\in \mathbb{R}^n$). In line 4, the minimum covariance determinant algorithm is used to compute the mean and the covariance matrix of samples in cluster $i$. In line 7, $\text{CDF}_{\chi^2_{d_1}}()$ is the cumulative distribution function of $\chi^2$ distribution with degree of freedom $d_1$.

---

**Algorithm 3:** FEATS Outlier Detection

---

**Input:** $X \in \mathbb{R}^{d \times n}$, $y$, $d_1$ and outlier_thres
**Output:** outlier_scores, outliers

1 $X_{\text{red}} \leftarrow \text{PCA}(X, d_1)$, where $X_{\text{red}} \in \mathbb{R}^{d_1 \times n}$ and $d_1 < d$
2 **for** $i = 1$ *to* $k$ **do**
3     Choose samples of $X_{\text{red}}$ belonging to cluster $i$ to get $X_i$, where $X_i \in \mathbb{R}^{d_1 \times n_i}$ and $n_i$ is the number of samples in cluster $i$
4     $(\mu_i, \Sigma_i) \leftarrow \text{MinCovDet}(X_i)$
5     **for** $j = 1$ *to* $n_i$ **do**
6         $D^2_{ij} \leftarrow (x_{ij} - \mu_i)^T \Sigma_i^{-1} (x_{ij} - \mu_i)$, where $x_{ij} \in X_i$ is the $j$-th sample in $X_i$
7 p_values $\leftarrow 1 - \text{CDF}_{\chi^2_{d_1}}(D^2)$, where $D^2 \in \mathbb{R}^n$ and $D^2 \sim \chi^2_{d_1}$
8 outlier_scores $\leftarrow -\log_{10}$(p_values)
9 outliers $\leftarrow$ outlier_scores $> -\log_{10}$(outlier_thres)

---
